# Investigating the Role of Glyoxalase 1 as a Therapeutic Target for Cocaine and Oxycodone Use Disorder

**DOI:** 10.1101/2024.12.23.630123

**Authors:** Elizabeth Alcantara, Michelle R. Doyle, Clara A. Ortez, Anne Ilustrisimo, Bloom Stromberg, Amanda M. Barkley-Levenson, Abraham A. Palmer

## Abstract

Methylglyoxal (MG) is an endogenously produced non-enzymatic side product of glycolysis that acts as a partial agonist at GABA_A_ receptors. MG that is metabolized by the enzyme glyoxalase-1 (GLO1). Inhibition of GLO1 increases methylglyoxal levels, and has been shown to modulate various behaviors, including decreasing seeking of cocaine-paired cues and ethanol consumption. The goal of these studies was to determine if GLO1 inhibition could alter cocaine-or oxycodone-induced locomotor activation and/or conditioned place preference (CPP) to cocaine or oxycodone. We used both pharmacological and genetic manipulations of GLO1 to address this question. Administration of the GLO1 inhibitor s-bromobenzylglutathione cyclopentyl diester (pBBG) did not alter the locomotor response to cocaine or oxycodone. Additionally, pBBG had no significant effect on place preference for cocaine or oxycodone. Genetic knockdown of *Glo1*, which is conceptually similar to pharmacological inhibition, did not have any significant effects on cocaine place preference, nor did *Glo1* overexpression affect locomotor response to cocaine. In summary, our results show that neither pharmacological nor genetic manipulations of GLO1 influence locomotor response or CPP to cocaine or oxycodone.

## 1. Introduction

Substance use disorders affected nearly 50 million people in the United States in 2023 (Substance Abuse and Mental Health Services Administration 2024). However, there are no pharmacotherapies approved by the United States Food and Drug Administration for cocaine use disorder, and only three approved compounds for opioid use disorder. Alterations in gammaaminobutyric acid (GABA) transmission have long been linked to substance use (Vaughan et al. 1997; Smith et al. 2009; Roberto et al. 2012; Kibaly et al. 2021). Additionally, GABA agonists, such as baclofen, have also been shown to decrease rewarding effects of cocaine and oxycodone in rodents (Roberts and Brebner 2000; Morris et al. 2008; van Zessen et al. 2012). Thus, there is interest in targeting the GABAergic system for treatment of substance use disorders.

A relatively novel strategy to target the GABAergic system involved increasing brain levels of methylglyoxal, an endogenous GABA_A_ receptor partial agonist (McMurray et al. 2017; de Guglielmo et al. 2018; Barkley-Levenson et al. 2018, 2021; Zhou et al. 2023). Since methylglyoxal is metabolized by the enzyme glyoxalase-1 (GLO1), administration of the the GLO1 pharmacological inhibitor s-bromobenzylglutathione cyclopentyl diester (pBBG) results in higher methylglyoxal levels (Distler et al. 2012). Recently, we reported a history of cocaine self-administration increases GABAergic transmission, especially in rats that exhibit high addiction-like behaviors (Zhou et al. 2023). These high addiction rats also show greater responding during a cue-induced reinstatement test. Importantly, both the increase in GABA transmission and the greater relapse-related effects can be reversed with administration of pBBG (Zhou et al. 2023). However, the role of GLO1 has not been investigated in other aspects of cocaine reward.

GLO1 has also been implicated in alcohol drinking using pBBG and *Glo1* knockdown or *Glo1* overexpressing mice (McMurray et al. 2017, 2018). Overexpression of *Glo1* increased drinking, and both the genetic knockdown of *Glo1* and pharmacological inhibition of GLO1 decreased drinking in the murine drinking-in-the-dark paradigm (McMurray et al. 2017). Importantly, both the genetic and pharmacological manipulations of GLO1 did not alter the locomotor-activating or sedative effects of alcohol nor increase withdrawal-induced seizures (Barkley-Levenson et al. 2018, 2021). In similar studies conducted in rats, we found pBBG administration reduced operant alcohol drinking in both dependent and nondependent rats (de Guglielmo et al. 2018).

Little work has been done with other substances; however, there is some evidence that pBBG administration can reduce fentanyl-induced locomotor activation in mice (Harp 2021) and an inbred strain with greater *Glo1* expression did not develop preference for fentanyl (Harp et al. 2022).

The goal of these studies was to extend the recent work to determine whether GLO1 can modulate other behaviors related to the rewarding effects of cocaine and oxycodone, and whether GLO1 may be a target for opioid use disorder. To do this, we evaluated the ability of pBBG to modulate the locomotor activation and rewarding properties of cocaine and oxycodone as measured by the conditioned place preference procedure. We also explored these behavioral drug responses in *Glo1* knockdown and *Glo1* overexpressing mice.

## 2. Materials and Methods

### 2.1. Animals

#### 2.1.1. Wildtype Mice and General Procedures

Adult male and female C57BL/6J mice (n=360) ordered from Jackson Laboratories at eight weeks of age and transgenic mice were obtained from our breeding colony (see below). Mice were allowed to acclimate following their arrival for a minimum of 72 hours before experimental testing began. Mice were housed two to five per cage and had access to food and water *ad libitum* and maintained on a 12h/12h light/dark cycle with lights on at 06:00. All procedures and tests were approved by the Institutional Animal Care and Use Committee at the University of California San Diego.

#### 2.1.2. Transgenic Mice

The transgenic *Glo1* overexpressing mouse line was previously generated in our lab on a FVB/NJ background (Distler et al. 2012). We used a BAC containing Glo1 with 35 copies of the gene, which increased GLO1 mRNA expression seventeen-fold (Distler et al. 2012). The *Glo1* knockdown mouse line was generated by Dr. Michael Brownlee in his lab at Albert Einstein College of Medicine, Bronx, NY on a C57BL/6J background and has a 45-65% reduction in *Glo1* gene expression (El-Osta et al. 2008). The transgenic mouse lines have been maintained by breeding heterozygous males to wildtype females obtained from Jackson Laboratories. Wildtype littermates were used as controls for the genetic manipulation experiments.

### 2.2. Drugs

Cocaine HCl (CAS # 53-21-4) and oxycodone HCl (CAS # 76-42-6) were purchased from Sigma Aldrich and dissolved in 0.9% sodium chloride and administered at doses of 10 and 1 mg/kg, respectively. Doses were selected because they have previously been shown to produce reliable CPP (Orsini et al. 2005; Niikura et al. 2013). The pBBG was synthesized at UCSD in the lab of D. Siegel (Skaggs School of Pharmacy and Pharmaceutical Sciences), and was dissolved in 8% DMSO (Sigma Aldrich), 18% Tween-80 (Sigma Aldrich), and 74% 0.9% sodium chloride and administered at doses of 12.5, 25, and 50 mg/kg. All drug treatments were administered intraperitoneally (i.p.) at a volume of 10 mL/kg body weight. All solutions were prepared fresh the day they were used, and vehicles for cocaine, oxycodone, and pBBG consisted of their respective vehicles.

### 2.3. Behavioral Experiments

#### 2.3.1. Conditioned Place Preference (CPP)

The conditioned place preference test was used to measure the motivational properties of cocaine (n=10-18/sex/pBBG dose) and oxycodone (n=8-24/sex/pBBG dose) in C57BL/6J mice, as well as in Glo1 knockdown mice (n=12-15/genotype). The CPP test was performed in a two-compartment box (42×42×31cm) with both visual and tactile cues. One compartment had white horizontal lines and a smooth flooring while the other had black vertical lines and a ridged floor. The two compartments were separated by a black divider with a dome shaped opening that allowed movement between the two compartments.

All mice underwent a pretest day (Day 0) where they received a saline injection and were placed in the box for 30 minutes with free access to both sides to measure any pre-existing compartment bias. Subsequently animals were placed in conditioning groups to have unconditioned bias by pseudo-randomly assigning animals to receive drug (10 mg/kg cocaine or 1 mg/kg oxycodone) on either the white or black compartment such that each group had an equal mean preference for both compartments. Mice were administered either saline or drug on conditioning days 1 and 3, and the opposite on days 2 and 4 and confined to the single side. On day 5, test day, mice were injected with pBBG (12.5, 25, 50 mg/kg) or vehicle (i.p.) 90 min prior to the test and saline immediately before being placed in the chamber. During the test, mice were allowed free access to both compartments for 30 minutes. For the experiment with Glo1 knockdown animals there was no pretreatment injection on day 5, but all other parameters remained the same. Position and distance traveled were measured by centroid body movement.

#### 1.3.2. Open Field Test (OFT)

OFT was conducted to measure whether manipulating GLO1 modified the locomotor response to cocaine or oxycodone. The studies were conducted in a clear box placed in a white sound attenuating cabinet equipped with a light and fan for background noise. Activity was monitored via photobeams in the VersaMax Legacy Open Field Chamber (Omnitech Electronics Inc.) and recorded through the Fusion program (v6.5 r1198 VersaMax Edition) for 30 minutes. Locomotor activity was measured as total distance traveled in centimeters by tracking centroid body movement using photobeams. Two different experimental paradigms were used for cocaine and oxycodone.

##### 2.3.2.1. Cocaine OFT

Locomotor response in an open field was used to determine whether pharmacological inhibition of GLO1 or genetic overexpression of *Glo1* could alter locomotor response to cocaine. C57BL/6J mice (n=15/sex/pBBG dose) were habituated to the chambers for two days prior to the OFT test. On the two habituation days, mice received i.p. saline injections immediately prior to the session. On the third test day, animals were pre-treated with pBBG (12.5, 25, or 50 mg/kg) or vehicle 90 minutes prior to the test, and then administered cocaine (10 mg/kg) immediately before being placed into the open field. The Glo1 overexpressing mice and their wildtype littermates (n=35-40/genotype) underwent the same paradigm, but did not receive injections of pBBG on the test day.

##### 2.3.2.2. Oxycodone OFT

Locomotor response to oxycodone was measured in mice that previously underwent a CPP experiment (see below), thus were not habituated to chambers as a part of the OFT paradigm. Mice (3-6/sex/pBBG dose) were used to test the interaction between oxycodone and pBBG. The mice were pre-treated with pBBG (12.5, 25, 50 mg/kg) or vehicle 150 min before the session, and then treated with oxycodone (1 mg/kg) or saline immediately before being placed into the OFT box.

### 2.4. Statistical Analysis

Results are shown as a mean ± standard error mean (SEM). Statistical analyses were performed using GraphPad Prism (Version 10). The CPP data was analyzed using a one sample t-test and a one-way ANOVA, with Dunnett’s post-hoc, when appropriate. Preference score was calculated by subtracting the time spent in the drug-paired compartment during the pre-test from the time spent in the drug-paired compartment on test day, where a positive score indicates preference for the drug-paired side. Locomotor activity data in the OFT was analyzed using a one-way ANOVA (cocaine), t-test (cocaine in *Glo1* overexpressing experiment), or two-way ANOVA (oxycodone). Factors such as sex, pass, and box # were included in the initial analysis in SPSS (Version 28) and removed after it was determined they had no significant main effects or interactions.

## 3. Results

### 3.1. Cocaine CPP

To evaluate whether pharmacological inhibition of GLO1 using pBBG could affect expression of place preference in C57BL/6J mice, vehicle or pBBG (12.5, 25, or 50 mg/kg) was administered before the test session (**Figure 1A**). A one sample t-test showed all groups expressed significant preference for the cocaine-paired side (ps ≤ 0.0063). However, a one-way ANOVA indicated no effect of pBBG dose (F (3, 110) = 0.75; p=0.53). There was no difference in locomotor activity between the four groups (F (3, 110) = 0.61; p=0.61) (data not shown).

**Figure 1.**
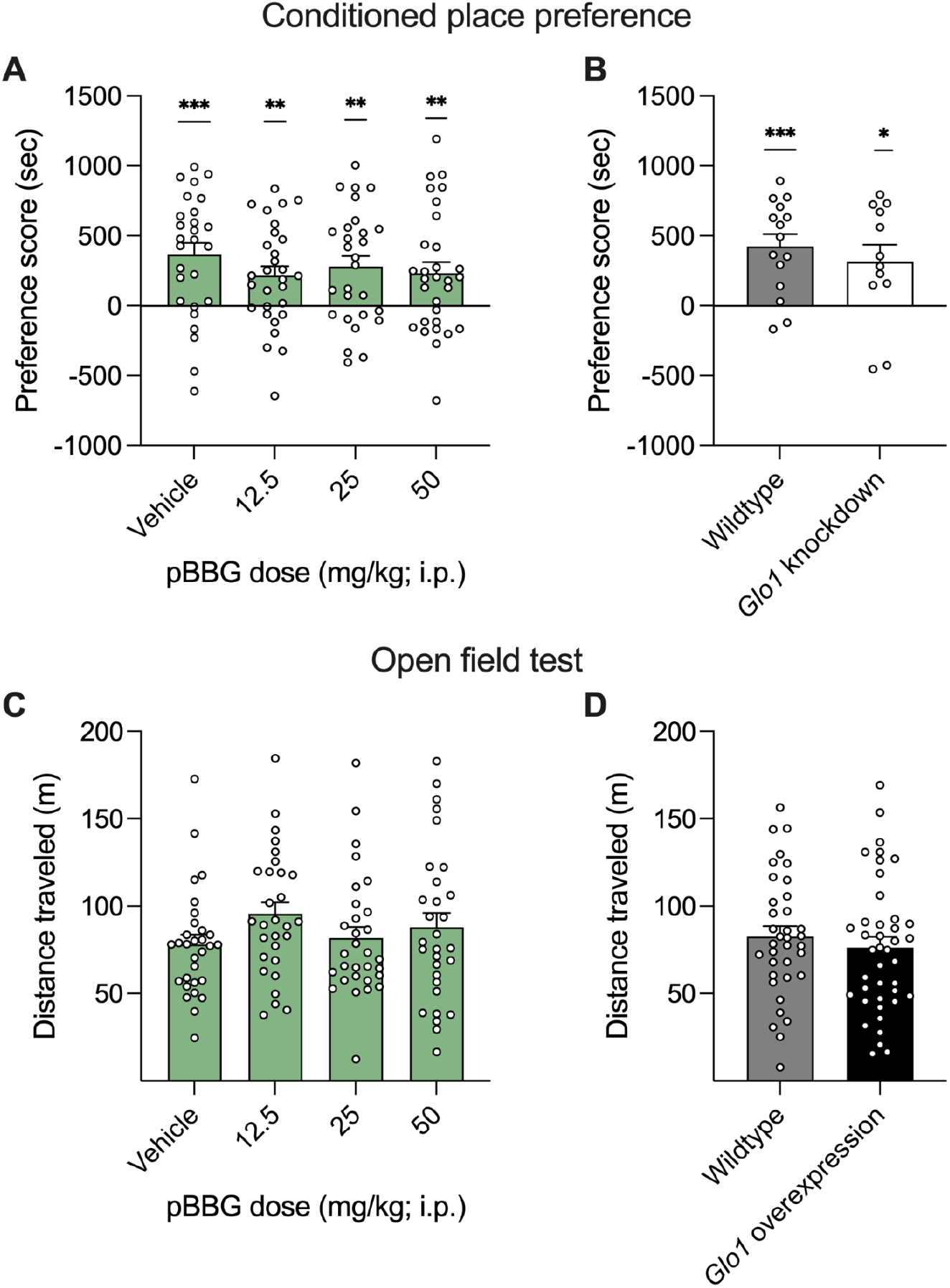
Conditioned place preference (CPP) and open field test (OFT) with cocaine. CPP produced by cocaine (10 mg/kg) in C57BL/6J mice treated with pBBG on the day of the test (A) or in Glo1 knockdown mice and their wildtype littermates (B). Total distance traveled in a 30-min test after administration of pBBG and 10 mg/kg cocaine in C57BL/6J mice (C) or Glo1 overexpressing mice and their wildtype littermates (D). Bars represent mean preference score (A, B) or distance traveled (C, D), error bars represent SEM, and points are individual mice. * represents p<0.05, ** represents p<0.01, *** represents p<0.001 when compared to 0 using a one sample t-test.

To determine whether genetic knockdown of *Glo1* affected cocaine place preference, cocaine CPP was conducted in *Glo1* knockdown mice and their wildtype littermates (**Figure 1B**). Both groups preferred the cocaine paired side (one sample t-test; ps ≤ 0.026); however there was no significant difference between the two groups (unpaired t-test; t=0.7564, df=25; p=0.46). Interestingly, the *Glo1* knockdown mice (54.5 ± 4.21 m) traveled greater distance compared to wildtype mice (41.0 ± 2.91 m) during the CPP test (t=2.711, df=25; p=0.012; data not shown).

### 3.2. Cocaine OFT

To determine whether pharmacological inhibition of GLO1 using pBBG affected cocaine-induced locomotor activity, pBBG (12.5, 25, or 50 mg/kg) or vehicle was administered prior to cocaine injection in an open field test (**Figure 1C**). pBBG administration did not alter total distance traveled (F (3, 115) = 1.277; p=0.2856). Similarly, genetic overexpression of Glo1 did not affect cocaine-induced locomotor activity (t=0.7302, df=73; p=0.47) (**Figure 1D**).

### 3.3. Oxycodone CPP

The effects of pharmacological inhibition of *Glo1* with pBBG (12.5, 25, or 50 mg/kg) or vehicle were measured on expression of oxycodone place preference (**Figure 2A)**. Although there was no difference between the groups (F (3, 87) = 0.46; p=0.71), all groups except the group that received 50 mg/kg of pBBG showed place preference (50 mg/kg: p=0.18; all other ps ≤ 0.049). There was a significant effect of pBBG dose on distance traveled (F (3, 87) = 5.12) where the group administered 50 mg/kg of pBBG (52.9 ± 3.11 m) had a significantly lower distance traveled compared to the vehicle group (34.5 ± 4.81 m) (p=0.0008) (data not shown).

**Figure 2.**
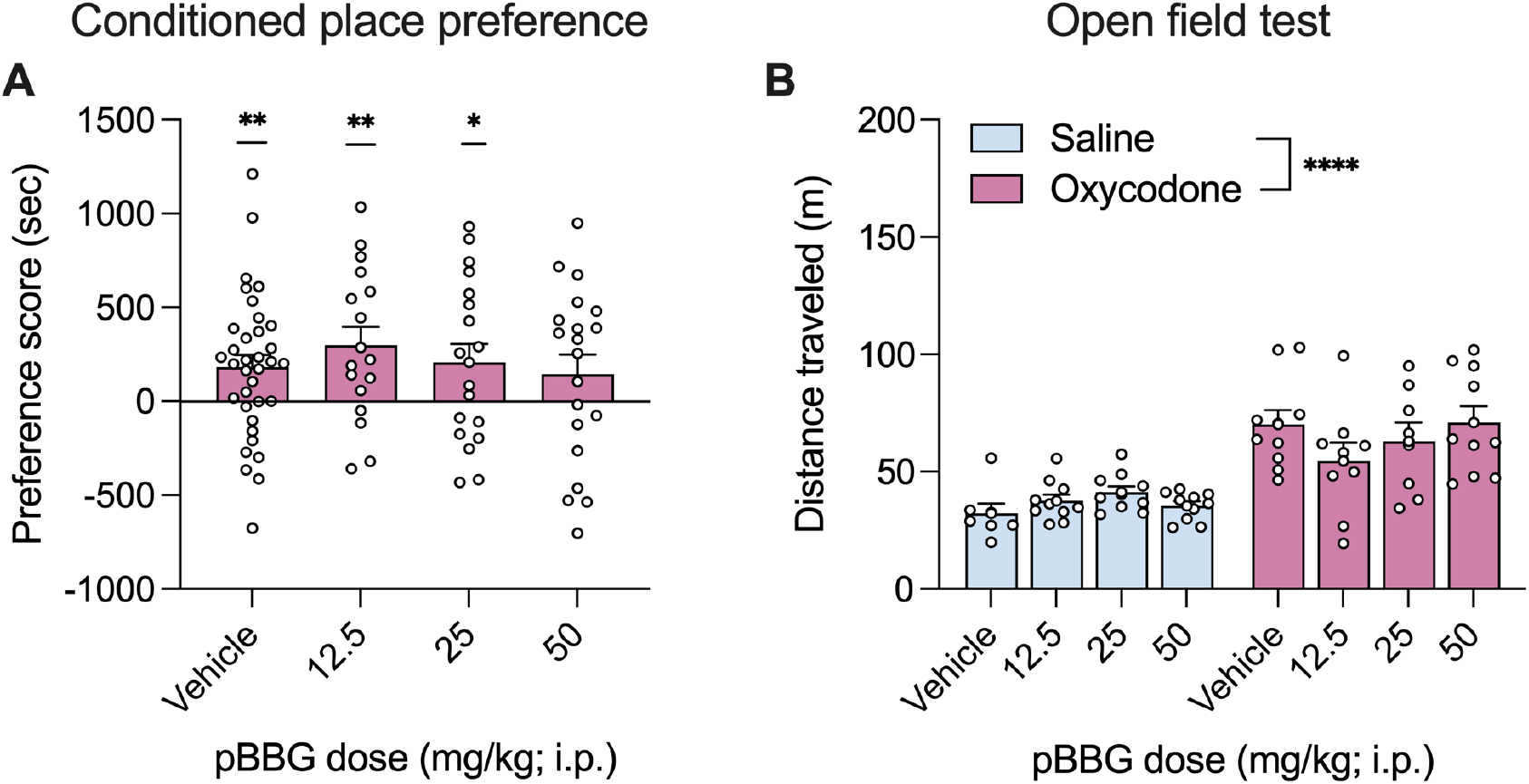
Conditioned place preference (CPP) and open field test (OFT) with oxycodone. CPP produced by oxycodone (1 mg/kg) in C57BL6/J mice treated with pBBG on the day of the test (A). Total distance traveled in a 30-min test after administration of pBBG and saline (blue) or 1 mg/kg oxycodone (pink) in C57BL6/J mice (B) Bars represent mean preference score (A) or distance traveled (B), error bars represent SEM, and points are individual mice. * represents p<0.05, ** represents p<0.01, *** represents p<0.001 when compared to 0 using a one sample t-test. **** indicated p<0.0001 for a main effect in a two-way ANOVA.

### 3.4. Oxycodone OFT

The effects of pBBG (12.5, 25, or 50 mg/kg) or vehicle on oxycodone-or saline-induced locomotor activity were measured in an open field test in the mice previously used in the CPP assay (**Figure 2B**). There was a significant effect of drug (oxycodone/saline: F (1, 66) = 52.25; p<0.0001) where mice administered oxycodone had greater distance traveled compared to those administered saline. However, there was no significant effect of pBBG (F (3, 66) = 0.6811; p=0.57) or significant interaction (F (3, 66) = 1.7; p=0.17).

## 4. Discussion

Novel pharmacotherapies are needed to treat substance use disorders and improve outcomes. The studies in this paper aimed to evaluate the utility of targeting GLO1 as a potential therapeutic for substance use disorders by using cocaine and oxycodone induced locomotor activity and place preference. We found that neither pharmacological inhibition of GLO1 nor genetic knockdown of *Glo1* were able to alter CPP for either cocaine or oxycodone. Similarly, neither pharmacological inhibition of GLO1 nor genetic overexpression of *Glo1* altered cocaine-or oxycodone-induced locomotor activation. Although these studies suggest pBBG did not affect locomotor activity or expression of CPP, both of which are sometimes considered to be proxies for reinforcing effects, other studies have suggested that pBBG is effective in rats that are dependent or have a long history of drug self-administration (Zhou et al. 2023).

Given that pBBG can reduce drug-seeking in cue-induced reinstatement tests in rats that previously self-administered cocaine (Zhou et al. 2023), it was somewhat surprising that pBBG did not alter expression of CPP in the cocaine assay. We found that neither pharmacological nor genetic manipulation of GLO1 was able to disrupt the rewarding properties of cocaine in the conditioned place preference test. In the previous study, pBBG was only effective in a subset of genetically diverse rats that were classified as having a high addiction index and not in those with a low addiction index (Zhou et al. 2023). In contrast, the mouse studies reported here were conducted in genetically identical C57BL/6J mice who had minimal drug exposure. Because our study used inbred mice, we were also unable to differentiate mice that may find cocaine more or less reinforcing to determine whether pBBG would have been effective in a subset of mice. Additionally, in the present study, mice only received two non-contingent injections of 10 mg/kg of cocaine, while the other study used rats with a long history of cocaine self-administration, which has shown to produce alterations in GABAergic signaling (Kallupi et al. 2013; Zhou et al. 2023). It seems likely that pBBG may only be effective at reducing rewarding properties of cocaine in animals with a long history of cocaine administration, thus future studies should pursue this line of research. There may also be important species differences since Zhou et al (2023)’s study used rats, whereas the present study used mice, in part to take advantage of the *Glo1* knockdown mouse lines.

Consistent with previous studies, these doses of pBBG did not have any effects on locomotion alone or altering cocaine-or oxycodone-induced locomotion in the open field test (McMurray et al. 2018; Barkley-Levenson et al. 2018). However, in the oxycodone CPP test, there was a reduction in overall locomotion in the mice administered the highest dose of pBBG. Given the negative results in other assays with the same dose of pBBG, both in this study and others, it is unclear what this interaction may indicate.

Since prior positive findings of the role of GLO1 on anxiety-like (Hovatta et al. 2005; Williams et al. 2009; Distler et al. 2012), and depression-like behavior (McMurray et al. 2018) and there is high comorbidity between mood disorders and substance use (Quello et al. 2005; Hartman et al. 2023), future studies should investigate the effectiveness of pBBG to attenuate anxiety-like behaviors and substance use. Increasing MG though GLO1 inhibition remains a possible therapeutic target and further studies should investigate its role in different reward circuitries.

## Acknowledgements

This work was supported by the National Institute on Alcohol Abuse and Alcoholism [R01 AA026281 (AAP), K99 AA027835 (AMB-L), and T32 AA007456 (MRD)]. The data that support the findings of this study are available from the corresponding author upon reasonable request. AAP holds a patent related to the use of GLO1 inhibitors (US20160038559, active). All other authors declare no competing interests.

## CRediT authorship contribution statement

**E. Alcantara**: Formal analysis, Investigation, Visualization, Writing -original draft. **M. R. Doyle**: Formal analysis, Visualization, Writing -original draft. **C. A. Ortez**: Investigation. **A. Ilustrisimo:** Investigation. **B. Stromberg**: Investigation. **A. M. Barkley-Levenson**: Conceptualization, Funding acquisition, Supervision, Writing -review & editing. **A. A. Palmer**: Conceptualization, Funding acquisition, Supervision, Writing -review & editing.

